# Radio-Immune Response Modelling for Spatially Fractionated Radiotherapy

**DOI:** 10.1101/2023.04.28.538767

**Authors:** Young-Bin Cho, Nara Yoon, John H. Suh, Scott G. Jacob

## Abstract

Radiation-induced cell death is a complex process influenced by physical, chemical and biological phenomena. Strong dose gradient may intensify the complexity and reportedly creates significantly more cell death known as bystander effect. Although consensus on the nature and the mechanism of the bystander effect were not yet made, the immune process presumably plays an important role in many aspects of the radiotherapy including the bystander effect. Immune response of host body and immune suppression of tumor cells are modelled with four compartments in this study; viable tumor cells, T cell lymphocytes, immune triggering cells, and doomed cells. The growth of tumor was analyzed in two distinctive modes of tumor status (immune limited and immune escape) and its bifurcation condition. Tumors in the immune limited mode can grow only up to a finite size, named as terminal tumor volume analytically calculated from the model. The dynamics of the tumor growth in the immune escape mode is much more complex than the tumors in the immune limited mode especially when the status of tumor is close to the bifurcation condition. Radiation can kill tumor cells not only by radiation damage but also by boosting immune reaction. The model demonstrated that the highly heterogeneous dose distribution in spatially fractionated radiotherapy (SFRT) can make a drastic difference in tumor cell killing compared to the homogeneous dose distribution. SFRT can not only enhance but also moderate the cell killing depending on the immune response triggered by many factors such as dose prescription parameters, tumor volume at the time of treatment and tumor characteristics. The model was applied to the lifted data of 67NR tumors on mice and a sarcoma patient treated multiple times over 1200 days for the treatment of tumor recurrence as a demonstration.

## 1 Introduction

Cells irradiated with sufficient energy of photons or charged particles are lethally damaged through physical, chemical and biological processes spanning time scales from pico-seconds to days (Zakaria et al. 2020; Tubiana, Dutreix, and Wambersie 1986). Irreparable cells from the damage will die and be removed from tissues with some time delay (Zhong and Chetty 2014; Lim et al. 2008). Radiotherapy of cancers has been evolved in a way to maximize the radiation damage to tumors respecting surrounding normal tissue tolerances. Considering tumor control probability sensitive to the minimum dose, extra efforts have been paid for the development of radiotherapy plans with homogeneous dose distribution conforming to targets. In recent decades, a new treatment delivery technique, FLASH (Chow et al. 2021; Yinghao Lv et al. 2022; Zakaria et al. 2020), using extremely high dose rate has reported a larger therapeutic gain than a standard radiotherapy due to the increased normal tissue tolerance. This finding highlighted novel treatments where extremely heterogeneous dose delivery in temporal domain can create. There is another way of delivering highly heterogeneous dose distributions with much longer history of practice namely spatially fractionated radiotherapy (SFRT) (M. Mohiuddin et al. 1999; Kanagavelu et al. 2014; Kaiser, M. M. Mohiuddin, and Jackson 2013). In this case heterogeneous distribution is in spatial domain and it is often reported to successfully treat extremely large tumors difficult to manage with a conventional radiotherapy. The mechanism of SFRT is presumably related to the tumoricidal effect on the untreated region as well as the treated part of tumors, known as the bystander effect(Yilmaz, Elmali, and Yazici 2019; Barsky et al. 2019; Wani et al. 2019; Bitran 2019; Hatten Jr et al. 2022; Ngwa et al. 2018; Leary et al. 2019). The improved therapeutic gain from such extreme spatio-temporal modulations in dose distribution may need deeper understanding of radiation interactions with cells in physical, chemical and biological levels.

Immune response is believed to play an important role in many aspects of the radiotherapy. Complete tumor regression is often observed with much less dose to kill all cancer cells suggesting radiation may trigger other tumoricidal mechanisms. Stone et. al.(Slone, Peters, and Milas 1979) reported that 1.67 time higher dose is required for tumor control of immuno-deficient mice compared to immuno-competent mice. Immune checkpoint inhibitors show outstanding results in a variety of cancers and combinations with radiotherapy are often desired for the synergistic effect for long lasting responses (Markovsky et al. 2019; Davidson et al. 2022; Vaage 1973). Thus it is crucial to develop a mechanistic model describing the dynamics of radio-immune response in order to understand the mechanism and efficiently develop clinical studies in this rapidly evolving area (Serre et al. 2016; Geng, Paganetti, and Grassberger 2017; Asperud 2020; Asperud 2020; Bekker et al. 2022).

To address this, Serre et al. 2016 developed a discrete-time numerical model of radio-immunotherapy in which radiation is prescribed with inhibitors of the PD1-PDL1 axis and/or CTLA4. This model offered an explanation for the reported biphasic relationship between the size of a tumor and its immunogenicity measured by the bystander effect. It also explained why discontinuing immunotherapy may result in tumor recurrence. In this model, tumor antigens are assumed to be released by tumor cells naturally at a constant rate and at higher rate after radiation exposure. Primary immune response is developed by immune effector cells (tumor infiltrating cytotoxic T lymphocytes (CTLs) or tumor associate macrophages, etc) proportionally generated by the volume of tumor antigens. Immune memory effect learned from the primary immune response was also modelled as a secondary immune response. It was demonstrated that the model can be successfully applied to simulate cancer immunotherapy and the synergy with radiotherapy. The Bystander effects previously reported in the experiment on mice (Vaage 1973) was explained using the model. Since the immune response is proportional to the irradiated volume in Serre’s model and partial irradiation in SFRT reportedly generates even larger immune response than full irradiation (Markovsky et al. 2019; Asperud et al. 2021; Asperud 2020; Kanagavelu et al. 2014; Kaiser, M. M. Mohiuddin, and Jackson 2013; M. Mohiuddin et al. 1999), Asperud et. al. (Asperud et al. 2021; Asperud 2020) developed a new numerical model with SFRT in mind. Instead of CTL generation directly from tumor antigens which are not killed by radiation in Serre’s model, Asperud assumed that CTLs need to be activated from a naturally inactive status. The rate of CTL activation was modelled as a function of irradiated tumor volume. The model also included time delay of cell clearance using a ‘doomed cell’ compartment for better agreement with *in vivo*. Damaged cells doomed to die by radiation and immunogenic damage are reportedly not removed from tissues instantly but can take days to be cleared (Zhong and Chetty 2014; Lim et al. 2008). The model was validated using previously published data on syngeneic xenografts (67NR breast carcinoma and Lewis lung carcinoma). This model, however, did not include the immune suppression effect of tumor cells and the immune boost by immunotherapy agents. In this study, we propose a new model to combine the strengths of two previous models; the effect of tumor volume and immunotherapy inhibitors from Serre’s model and the mechanism of active/inactive CTL and doomed cells in Asperude’s.

The rest of the manuscript is structured as follows. The first section will describe the development of the mathematical tumor model for radio-immune response. In the following sections, analytic solutions of the tumor growth are presented in distinctive boundary conditions distinguished by the bifurcation threshold; immune limited (trapped) *vs* immune escape. The pattern of tumor growth changes drastically around this bifurcation threshold. Mouse experiment on syngeneic xenografts (67NR breast carcinoma) by Asperud (Asperud et al. 2021) and a single case of patient data were used for an application of the developed model.

## 2 Methods

### 2.1 Radio-immune response model

The tumor model in this study consists of four compartments: the volume of viable cancer cells *T*_*n*_, doomed cells *D*_*n*_ which represents dead cells to be removed from the host body with some time delay after lethal damage, active cytotoxic T lymphocytes (CTLs) *L*_*n*_ responsible for the anti-tumor immune response and the density of immune triggering cells *A*_*n*_ converting naive CTLs to be active on attacking tumor cells. The measure of time *n* is expressed as days for all compartments. The schematic is shown in **Fig. 1**.

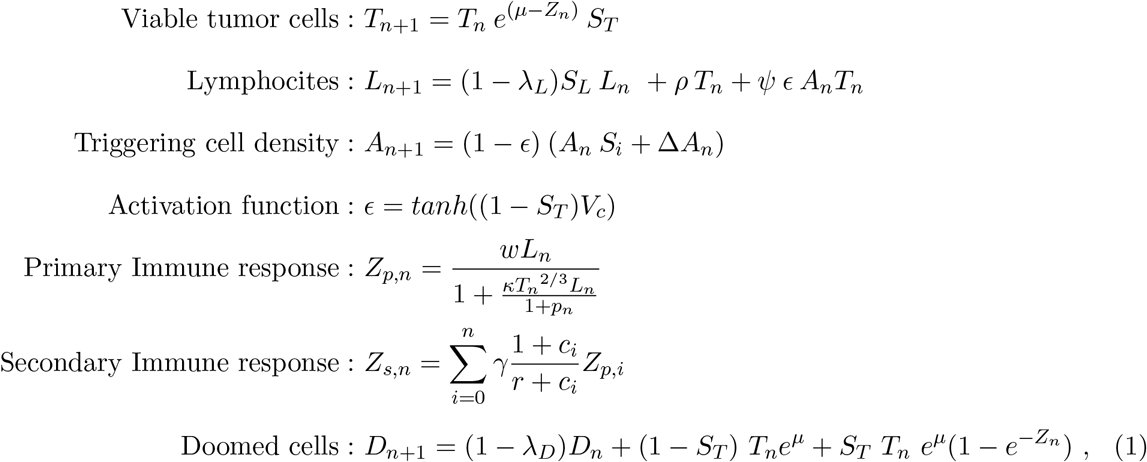

where total immune response is the sum of the primary and secondary immune effect *Z*_*n*_ = *Z*_*p,n*_ + *Z*_*s,n*_. Parameters and numerical values used in this study are summarized in **Table 1**.

**Figure 1:**
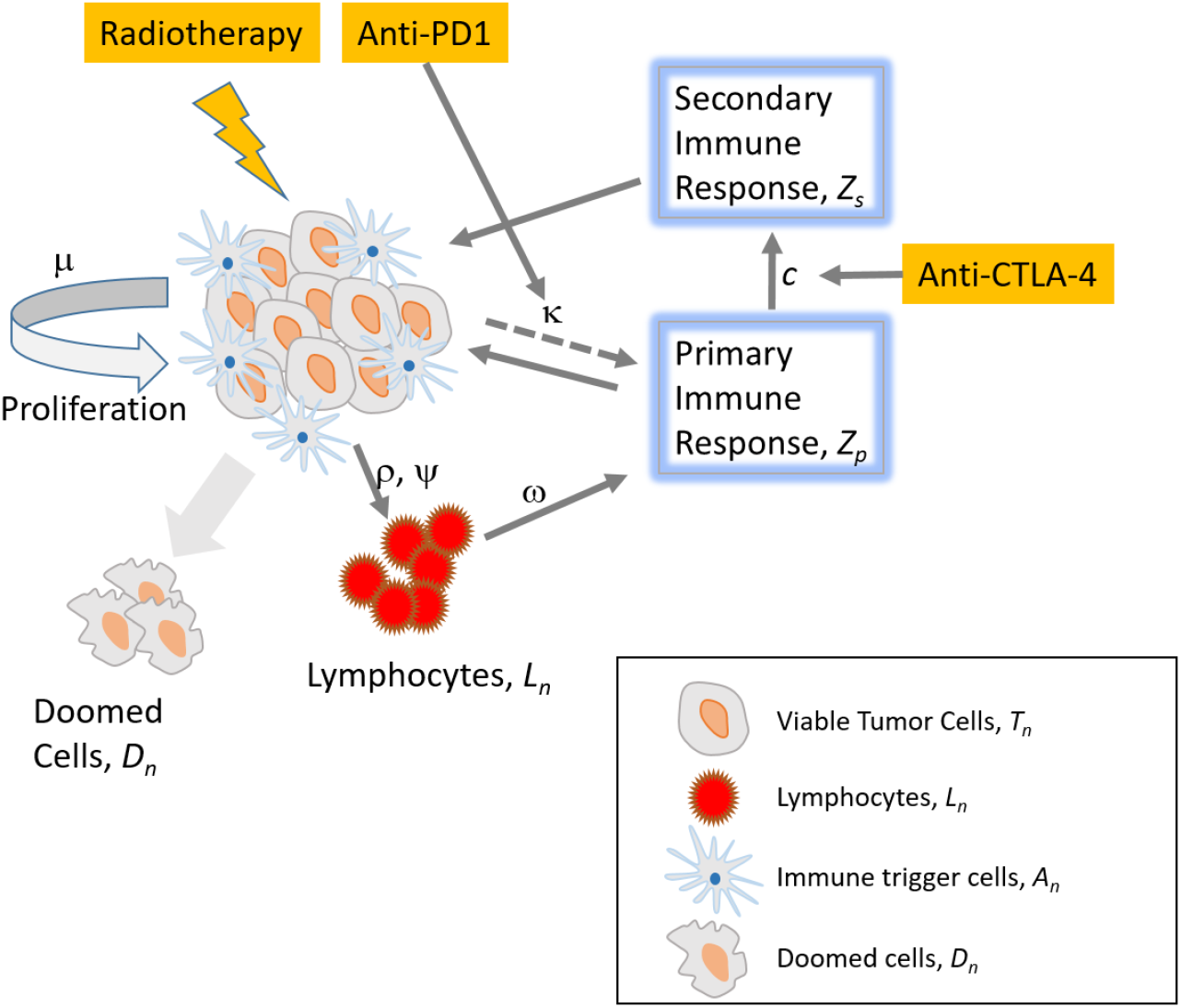
Schematic of the tumor model. Viable tumor cells proliferate exponentially with growth rate of *μ*. Tumor cells lethally damaged by either radiation or immune response turn into doomed cells to be cleared from the body. Lymphocytes are activated by various mechanisms including tumor antigen release in natural and radiation damaged tumor cells. Lymphocytes attack tumor cells by immune response and tumors fight back through an immune suppression mechanism. Immunotherapy can affect the immune responses.

**Table 1:**
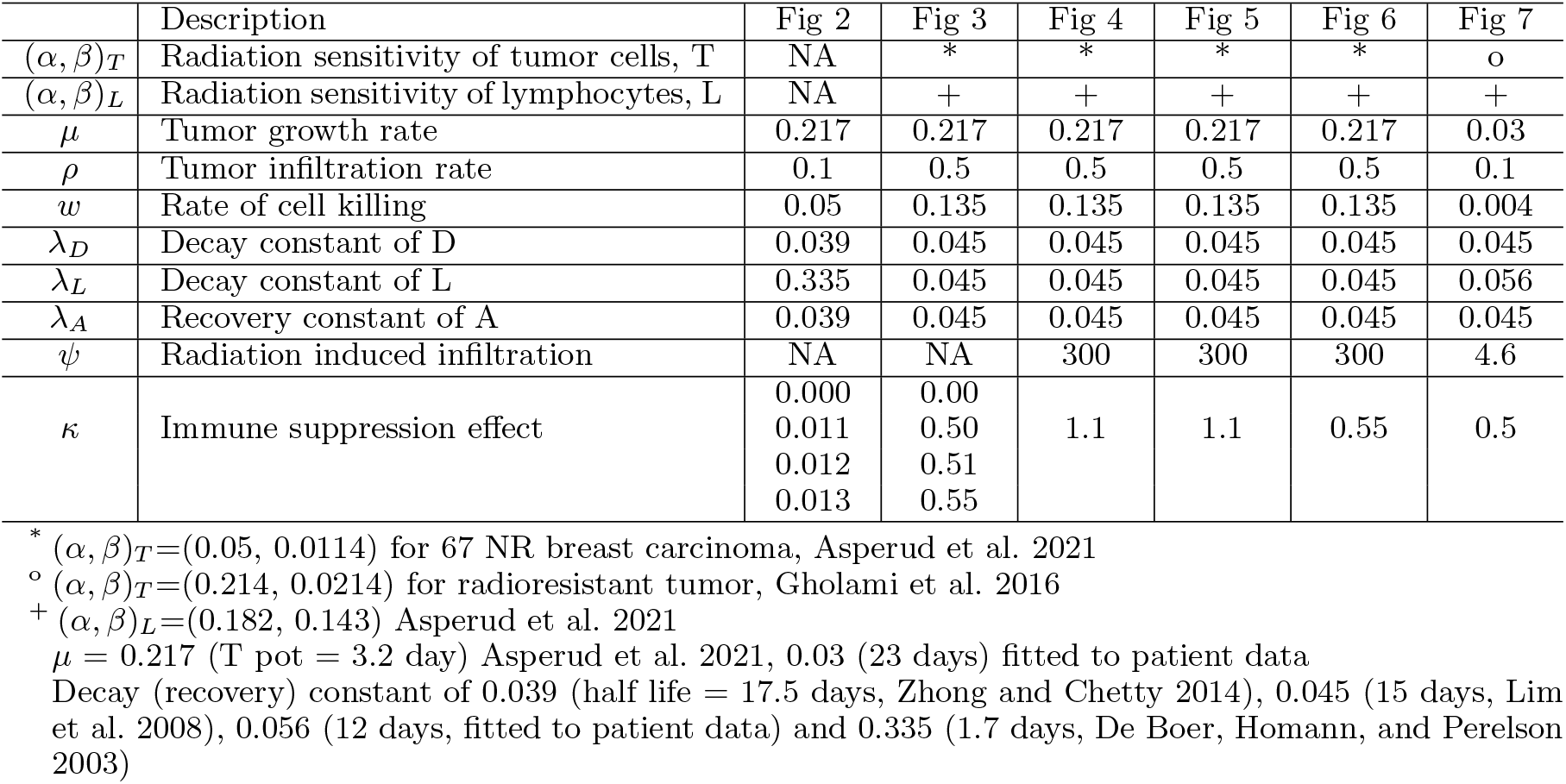
Summary of model parameters.

#### Viable Tumor Cells

The growth rate of tumor *μ* is a small positive constant. In the absence of immune response (*Z*_*n*_ = 0), the tumor will grow exponentially. This simplification has been adopted previously for the experimental results on small size tumors (Asperud 2020; Serre et al. 2016). Although other sophisticated tumor dynamics can be considered such as the Gompertz model (Gompertz 1833), the simplification should be adequate for larger tumors since the improvement of the modelling accuracy from these advanced models is based on the consideration of tumor volume effect that might be modelled in the immune dynamics (Gerlee 2013). The immune response *Z*_*n*_ is assumed acting as an anti-tumor growth factor which slows down the tumor growth or even shrinks the tumor volume depending on its intensity. The surviving fraction (*S*) from irradiation is based on the generalized linear-quadratic (LQ) model defined as 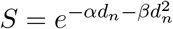 where (*α, β*) is radiation sensitivity and *d*_*n*_ is radiation dose. The surviving fraction for tumor, CTLs and immune triggering cells are expressed with *S*_*T*_, *S*_*L*_ and *S*_*i*_, respectively.

#### Lymphocytes

There is a growing body of evidence indicating radiation-mediated T-cell priming through the activation of host immunity (Lee et al. 2009; Twyman-Saint Victor et al. 2015; Walle et al. 2018; Davidson et al. 2022). Tumor antigens released either from tumors in natural condition or from radiation damaged tumors lead to the priming of tumor antigen specific T-cells. A positive feedback loop is often established by another antigens released from the tumors attacked by the T-cells. Radiation, therefore, may provoke dendrite cell mediated tumor antigen specific T-cell priming, transforming a tumor into an *in situ* vaccine (or create vaccine instead) (Walle et al. 2018). The finite life span of CTLs is modelled by the decay constant 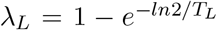, where *T*_*L*_ is the half life of CTLs and by radiation surviving fraction *S*_*L*_. The naive CTLs are activated in natural conditions proportional to the tumor volume. This is modelled by tumor infiltration rate, *ρ*. The CTL activation can be further amplified proportionally (at the rate of *ψ*) to the radiation-damaged tumor cells, *ϵT*_*n*_ and the amount of immune triggering cells (such as dendrite cells attracted by ATP released from tumors), *A*_*n*_. The immune triggering cells is modelled to be 1.0 in fully saturated conditions before radiation and can be reduced by radiation damage determined by surviving fraction *S*_*i*_. The rate of CTL activation *ϵ* depends on the relative volume of radiation-damaged cancer cells. We use the hyperbolic tangent to limit the range between 0 and 1. The CTL activation from radiation-damaged tumors is thus completed by *ψϵA*_*n*_*T*_*n*_.

#### Triggering Cell Density

Immune triggering cell density *A*_*n*_ is reduced at the rate of *ϵ* after the activation of CTLs and can be damaged by radiation. Surviving fraction of immune triggering cells is denoted by *S*_*i*_. For long-term analysis such as re-treatment cases taking months or years, *A*_*n*_ can be further assumed to be slowly saturated back after the radiation damage Δ*A*_*n*_ = (1 −*A*_*n*_)*λ*_*A*_, where *λ*_*A*_ is the recovery constant similarly defined as *λ*_*L*_.

#### Immune Response

The primary anti-tumor immune response is assumed proportional to the amount of CTLs with the rate of cell killing, *w* when there is no immune suppression effect, *κ* = 0. It is known that the immune response is tempered by tumor cells proportional to the surface area of tumor volume and the amount of CTLs, *L*_*n*_. Thus immune suppression effect is modelled by 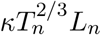 (Serre et al. 2016). Immunotherapy can modify the immune suppressive capability of cancer cells. In our primary immune response equation, *p*_*n*_ is the effect of PD1-PDL1 immunotherapy to control the tumor’s immune suppression. *Z*_*s*_ is the secondary immune response or memory effect learnt from primary immune reaction, *γ* is the sensitivity factor for *Z*_*s*_, *r* is the normalization factor and *c*_*n*_ is the effect of CTLA4 immunotherapy. A detailed explanation on immunotherapy effect can be found in the literature (Serre et al. 2016).

#### Doomed Cells

After lethal damage, it can take up to 1-2 weeks to clear these cells from the body. Therefore *in vivo* tumor volume measurement may not correlate well with viable tumor cells. Sum of viable tumor cells and doomed cells may be related better with the measurement (Asperud et al. 2021). The volume of damaged CTLs and immune triggering cells are not included in the doomed cell formulation in this analytic model for simplicity. Decay constant of doomed cell *λ*_*D*_ is expressed in a similar way as *λ*_*L*_. The amount of CTLs in the formulation can be reduced asymptotically when *ϵ* is small.

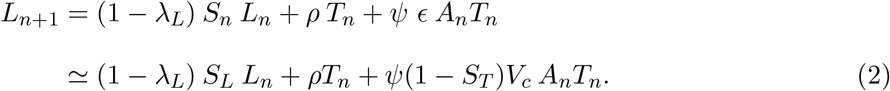

The delay between radiation damage and the death of tumor cells is modelled using an aggregated survival probability model (Serre et al. 2016). In this study, the same mean of 3 days and the standard deviation of 1.5 days were used in a log-normal density probability function which translates 95% of lethally damaged cells died in 6 days after radiation.

### 2.2 Boundary behavior

There exist distinct phases dictated by boundary conditions in the proposed numerical model. Some of the fundamental phases are reviewed in this section in order to understand the mechanism of radio-immune dynamics. Although the model is capable of considering immunotherapy drugs, for simplicity, we do not consider these in the boundary behavior analysis. The secondary immune response is also neglected in the analysis and simulations.

#### 2.2.1 Case 1: Immune limited

One of the simple boundary conditions would be tumors with the lack of secondary immune capability (*γ* = 0), inability of immune suppression effect (*κ* = 0), no use of immunotherapy drugs (*p* = 0) and no radiation (all *S* = 1). In this case, the tumor will start to grow exponentially at the initial phase and the growth rate slows until it stops growing completely at the equilibrium condition (*Z*_*n*_ = *μ*). The anti-tumor immune response, *Z*_*n*_ is linearly increasing with the tumor volume in this condition. The terminal viable tumor volume, *T*_∞_ in the equilibrium condition can be found with *L*_*n*+1_ ≃ *L*_*n*_ and *S*_*L*_ = *S*_*T*_ = 1 in Eq. (2).

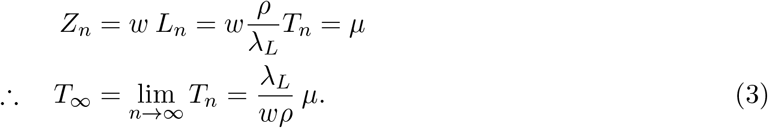

The amount of doomed cells in the equilibrium condition can be found similarly as shown below.

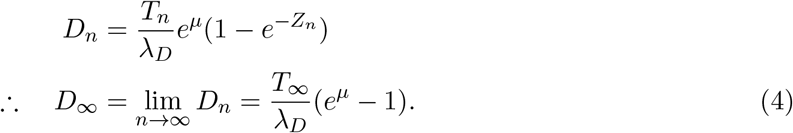

This equilibrium condition is an example of an immune limited, likely pre-malignant tumor that requires immune escape to grow further (one of the hallmarks of cancer, R. Weinberg and Hanahan 2000; Hanahan and Robert A Weinberg 2011).

#### 2.2.2 Case 2: Immune escape and bifurcation condition

When immune suppression becomes larger than a certain threshold 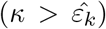 with all other conditions the same as Case 1, tumors can break the equilibrium and keep growing (immune escape). In this condition, the volume of active CTLs also keep growing with the tumor volume. The bifurcation threshold to break the equilibrium, 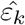 can be found as shown below,

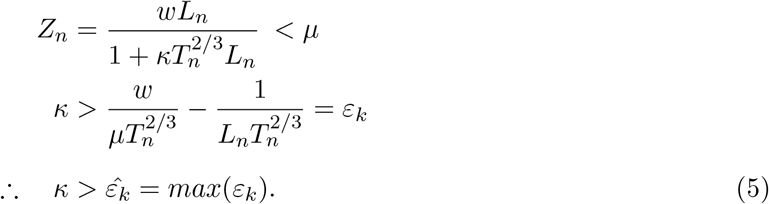

The maximum *ε*_*k*_ is found at 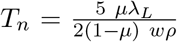 or 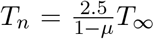 from the solution of 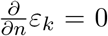 with the approximation of **Eq**. (2) such that 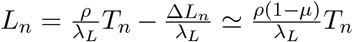 for Δ*L*_*n*_ ≃ *ρ*Δ*T*_*n*_ = *ρμT*_*n*_ in early phase of growth where tumor growth dominates the immune reaction and breaks equilibrium condition. The bifurcation threshold therefore can be found as,

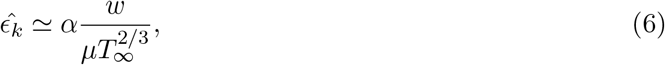

where 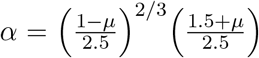. As will be discussed later, **Eq**. (6) is an approximated solution for the bifurcation threshold, it agrees well for stably transient cases but tends to over-estimate for unstable transient cases where both immune response and immune suppression are very strong. In other words, tumors with highly unstable transient dynamics may escape the equilibrium condition earlier than the approximated solution of the bifurcation threshold in **Eq**. (6).

#### 2.2.3 Case 3: Generalized expression

Immune effect *Z*_*n*_ can be rewritten by substituting *κ* with 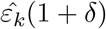, where *δ* ≥ −1. With this representation, Case 1 and 2 can be considered as a special case of the general representation with *δ* = −1 for Case 1 and *δ* = 0 for Case 2. Tumors can be in one of the conditions below,

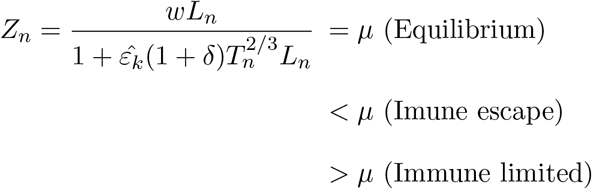

The equation above can be represented by substituting *κ* with 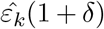 using **Eq**. (6). Tumors at the condition of immune limited is written below,

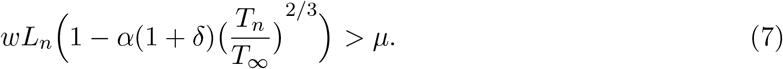

The left side of Eq. (7) can be considered as a general form of the immune effect, *Z*_*n*_:

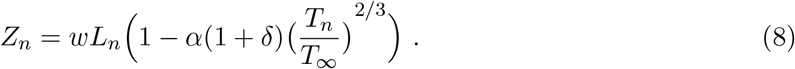

As previously described, **Eq**. (8) is reduced to Case 1 and 2 with *δ* = −1 and *δ* = 0, respectively. For tumor with immune escape capability, therapeutic intervention is needed to control the tumor volume. Radiotherapy can kill tumor cells not only by radiation damage but also by boosting CTL generation through the process expressed by the term of *ψϵA*_*n*_*T*_*n*_ in **Eq**. (2). To keep the tumor volume under control, it is necessary to have the parenthesis in **Eq**. (8) a positive value. The condition is translated that the radiation treatment should start early enough before the tumor volume reaches the critical tumor volume shown below,

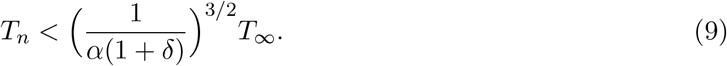

The generalized terminal viable tumor volume, *T*_∞,*δ*_ can be found from the solution of *Z*_*n*_ = *μ* in **Eq**. (8) with Taylor expansion of *T*_*n*_*/T*_∞_,

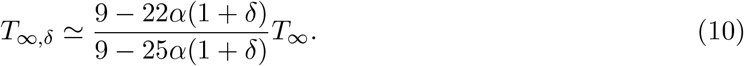

The terminal viable tumor volume in Case 1, *T*_∞_ is a special case with *δ* = −1.

## 3 Results

We analyzed our novel model of radio-immuno response and found two distinctive boundary conditions as described in the previous sections. In **Fig. 2 (A) and (B)** we show the terminal tumor volume with potential doubling time of 3.2 days (67NR mice data, Markovsky et al. 2019) for an illustration of the **Case 1 (***κ* = 0.0). The primary immune effect and the decay constant of CTL are modeled with *ρ* = 0.1, *w*=0.05, *λ*_*D*_=0.039 (half life of 17.5 days) and *λ*_*L*_ = 0.335 (half life of 1.7 days), respectively (Asperud et al. 2021; Asperud 2020). Full model parameters can be found in **Table 1**. The immune effect *Z*_*n*_ is increasing with tumor volume and reaches the equilibrium status equal to the tumor growth rate, *μ* (**Fig. 2 A**). The tumor volume *T*_*n*_ and the doomed cell volume *D*_*n*_ converge well to the estimated terminal volume of 14.5 cc and 90.7 cc, respectively as shown in horizontal dash lines in **Fig. 2 (B)** calculated with **Eq**. (3) and (4). The dynamics of the immune effect, *Z*_*n*_ looks like that of the viable tumor volume but with some time lag about the half life of lymphocyte, *λ*_*L*_. An exponential tumor growth is shown in the dotted line as a reference when no primary immune reaction exists in the cells or immune suppressed subjects are considered. Tumors with immune suppression capability (*κ >* 0) can grow larger than those without the capability as shown in **Fig. 2 (C) and (D)**. As immune suppression is larger, immune effect suffers and is eventually suppressed below the tumor growth rate, *μ*. This leads to catastrophic tumor growth, called bifurcation condition as described in **Eq**. (6). The terminal tumor volume at the equilibrium, *T*_∞,*δ*_ increases with immune suppression effect, *κ* (or equivalently *δ*) as expected in **Eq**. (10) and as shown in **Fig. 2 (D)**. The maximum viable tumor volume in the immune limited mode is 27.9 cc well within the bifurcation limit of 30.9 cc calculated from **Eq**. (9).

**Figure 2:**
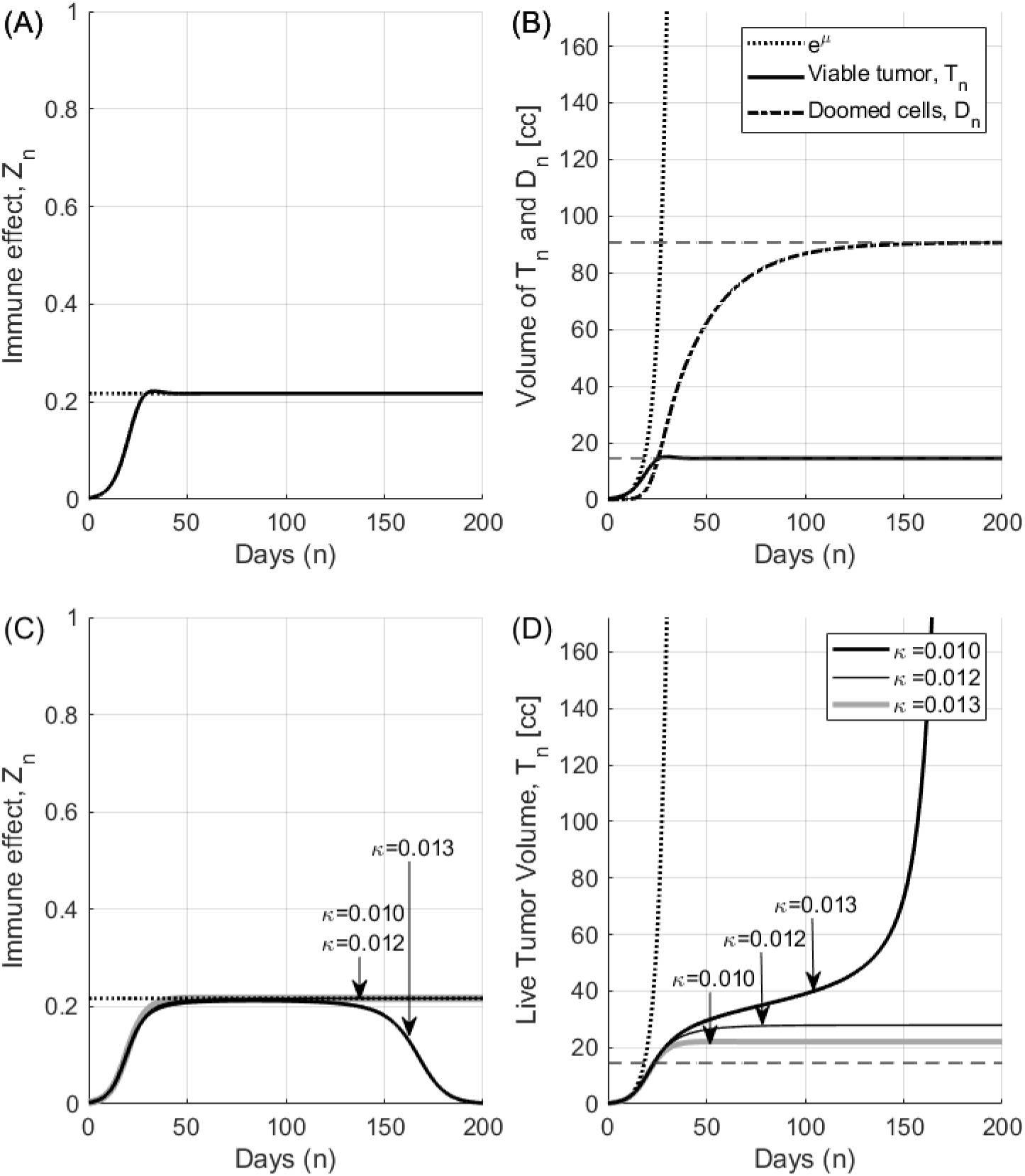
Dynamics of tumor immune effect, terminal viable tumor volume (Case 1) and the bifurcation of tumor growth (Case 2). It is assumed that *μ*=0.217 (*T*_*pot*_ = 3.2 days), *ρ* = 0.1, *w*=0.05, *λ*_*D*_ =0.039 (half life of 17.5 days) and *λ*_*L*_ = 0.335 (half life of 1.7 days). Detailed model parameters can be found in **Table 1. (A)** With no immune suppression (*κ* = 0), immune effect *Z*_*n*_ reaches the equilibrium status equal to the tumor growth rate, *μ* depicted with a dashed horizontal line. **(B)** Viable tumor cells *T* and doomed cells *D* converge to the estimated terminal volume of 14.5 cc and 90.7 cc as shown in horizontal dashed lines calculated with **Eq**. (3) and (4). Exponential tumor growth is shown in the dotted line as a reference in case of no immune effect or complete immune suppressed subjects. **(C)** Immune effect, *Z*_*n*_ starts suffering from immune suppression (*κ >* 0.0). It is hindered completely when *κ* is larger than the bifurcation threshold (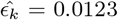 in this example). **(D)** The terminal viable tumor volume, *T*_∞,*δ*_ is increasing with *κ* as shown in **Eq**. (10). The horizontal broken line shows the terminal viable tumor volume at *κ* = 0 in Case 1 (*T*_∞_) for comparison. Tumor can grow out of the equilibrium condition and enters into the immune escape mode when *κ* is larger than the bifurcation threshold.

Tumors with stronger immune reactions can result in a smaller terminal tumor volume but the change in tumor volume becomes more dynamic. As an example, tumors with the same doubling time of 3.2 days but with much stronger immune response of *ρ* = 0.5, *w*=0.05, *λ*_*D*_=0.045 (half life of 15 days) and *λ*_*L*_ = 0.045 (half life of 15 days) are considered in **Fig. 3**. Larger infiltration rate *ρ* and longer half life of CTLs *λ*_*L*_ represent a stronger immune reaction. The terminal viable tumor volume *T*_∞_ is 0.15 cc much smaller than 14.5 cc in **Fig. 2**. The immune effect *Z* reaches the equilibrium status with dynamics akin to an attenuated oscillation due to the competing actions of tumor growth (*μ*) and immune reaction (*ρ* and *w*) for tumor without immune suppression capability, *κ* = 0. The shape of the immune effect, *Z* looks like the shape of the viable tumor volume again with some time lag about the half life of lymphocyte, *λ*_*L*_. This oscillating behavior gets amplified with increasing immune suppression effect, *κ* and the tumor growth finally breaks out of the equilibrium. As shown in **Fig. 3 (B)**, immune escape occurs at *κ* = 0.51 which is smaller than the bifurcation threshold, 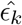 of 0.72 calculated from **Eq. 6**. The maximum viable tumor volume before bifurcation is 1.02 cc well within the limit of 1.39 cc from **Eq**. (9). The early escape is due to the dynamic characteristics of tumor volume overshoot in this condition. Assumptions used in the derivation of **Eq**. (6) such as steady change of status are responsible to this discrepancy. Such complex tumor dynamics when host body’s strong immune system vigorously fights against tumor’s immune suppression becomes more complex when radiation is involved. The effect of immune response in partial or full irradiation is simulated in **(C)-(F)**. Radiosensitivity (*α, β*) of tumor and CTLs is assumed to be (0.05, 0.0114) and (0.182, 0.143), respectively in the unit of [*Gy*^−1^, *Gy*^−2^] with *ψ* = 300 and *κ* = 0.55. Note that *κ* is well within early immune escape condition with some margin but still less than the bifurcation threshold of 0.72. Partial irradiation (50% tumor volume) with 20 Gy triggers larger immune reaction **(C)** at early phase compared to full irradiation **(E)** with 20 Gy. As a result, the time to tumor re-growth is also larger for partial irradiation in **(D) and (F)**. When adequate amount of dose (1 Gy ≤ dose ≤ 10 Gy in single fraction) is given to the whole tumor volume (full irradiation) in **(E) and (F)**, the immune effect becomes similar to the level of tumor growth factor. As a result tumor volume can stay at the equilibrium condition long period time with some visible oscillation. Since tumor cannot make early immune escape, it can be considered as arrested in the immune limited condition. Partial volume irradiation does not show such arrest event.

**Figure 3:**
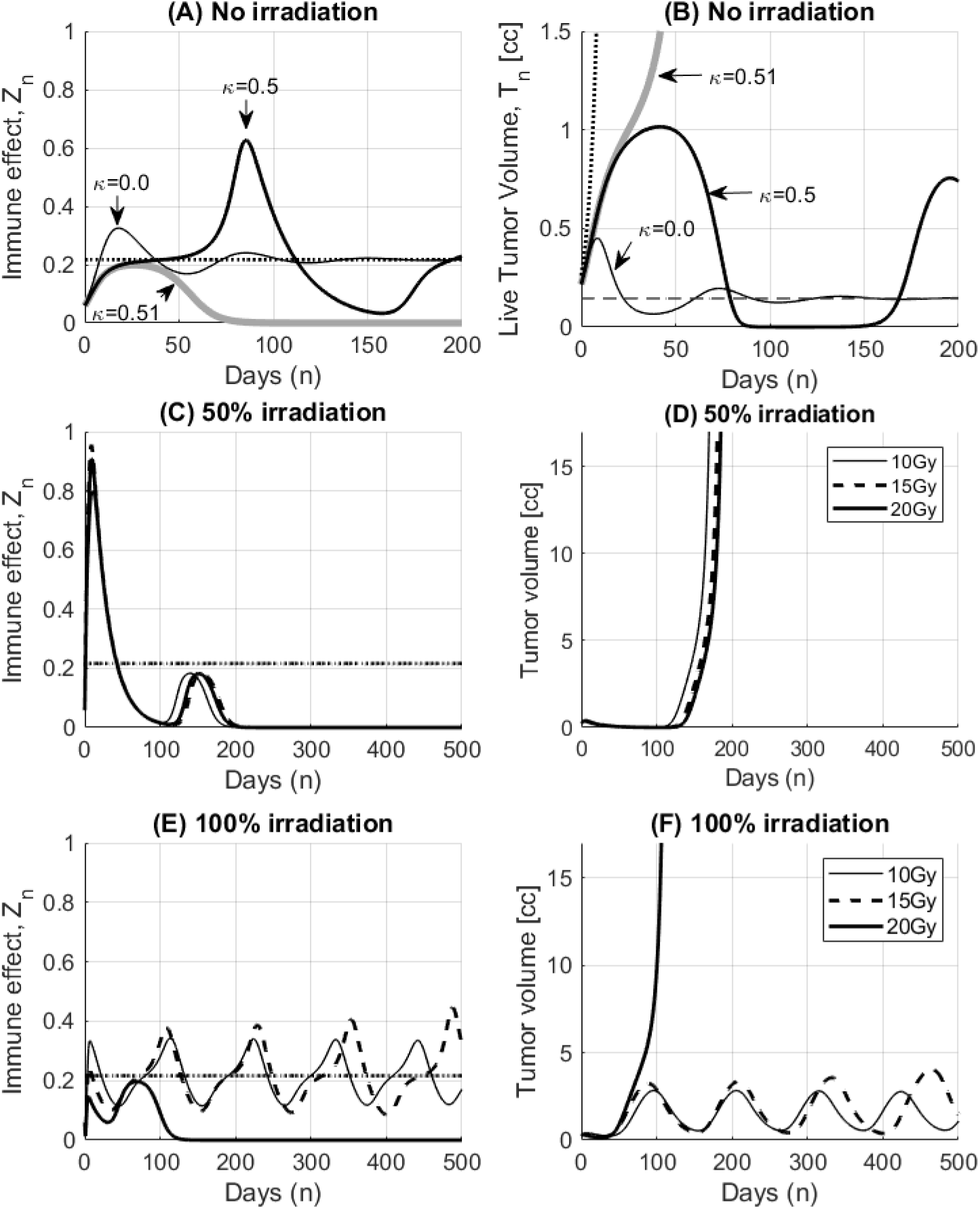
Dynamics of tumor immune effect and the bifurcation of tumor growth. In this example, the same tumor doubling time but much stronger immune reaction (larger *μ, w* and smaller *λ*_*L*_, *λ*_*D*_) is assumed. **(A)** Immune effect shows unstable dynamics over time. The magnitude of instability increases with the immune suppression effect, *κ*. When *κ* becomes larger than certain level (0.51 in this case) however, the immune effect is suppressed most of the time and the tumor can change its mode to the immune escape (continuous tumor growth). **(B)** The terminal tumor volume *T*_∞_ is 0.15 cc shown in the horizontal dash line which is only about 10% of **Case 1** (**Fig. 2**) due to the stronger immune reaction. The bifurcation occurs at *κ* = 0.51 (early immune escape), smaller than the solution of 0.72 from **Eq. 6**. In **(C)-(F)** *κ* is set to 0.55 which is an early immune escape condition with some margin but still less than the bifurcation threshold of 0.72. Partial irradiation with 20 Gy triggers larger immune reaction **(C)** compared to full irradiation **(E)** with 20 Gy. When adequate amount of dose (1 Gy ≤ dose ≤ 10 Gy in single fraction) is given to the whole tumor volume in **(E) and (F)**, the immune effect becomes similar to the level of tumor growth factor (arrested). Partial volume irradiation does not show such arrest event. Tumor volume in vertical axis represents *T*_*n*_ + *D*_*n*_.

The effect of immune response in partial or full irradiation on 67NR tumors is simulated in **Fig. 4**. All other parameters are the same as in the previous simulation **Fig. 3** except *κ* = 1.1. Total tumor volume (sum of the viable tumor volume, *T*_*n*_ and the doomed cell volume, *D*_*n*_) is compared with the measurement data originally published by Markovsky et al and digitally lifted later by Asperud et al. 10 Gy of radiation is delivered in a single fraction on either 50% (partial) or 100% (full volume) of the tumor volume. Due to the larger radiation damages, the tumor volume is reduced faster in the full irradiation case especially in early phase (*<* 20 days) as shown in **Fig. 4 (B)**. However the less damaged CTLs from the partial irradiation trigger immune reaction in a higher level for longer period (**Fig. 4 (A)**). As a result, the tumor volume is controlled better at later phase (*>* 20 days) with the partial irradiation as shown in **Fig. 4 (B)**. Simulation result is extrapolated to the range of 150 days in **Fig. 4 (C)**. Although the extrapolated result cannot be validated since the measurement is available only up to 30 days in the original study, the hypothetical behavior of the tumor growth under partial and full irradiation can be reviewed with the model. With much more activated CTLs, the time delay of tumor re-growth is expected greater for the partial volume irradiation.

**Figure 4:**
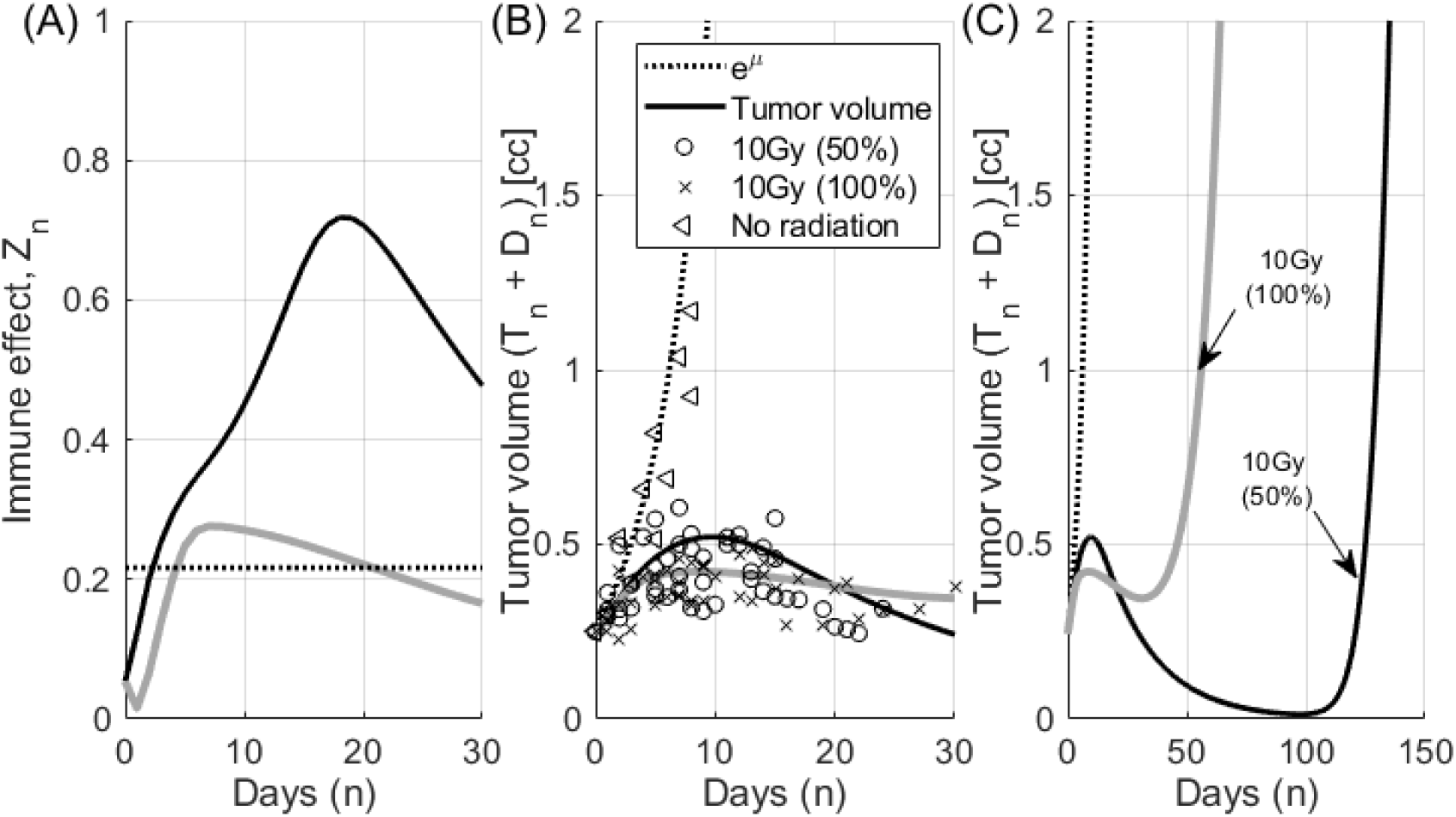
Radio-immune response of 67NR tumors (Markovsky et al, Asperud et al). Tumors (mean volume of 0.25 cc) were irradiated with 10 Gy either at 50% (in gray) or 100% coverage (in black) in the experiments. Model parameters including radiation sensitivity are summarized in Table 1. The *S*_*i*_ is assumed to be the same as *S*_*L*_. (A) Immune effect is larger for the partial irradiation due to the less radiation damaged immune triggering cells and CTLs. (B) Total tumor volume (*T*_*n*_ + *D*_*n*_) in solid black line is compared with the measurements. The measurement points were lifted digitally (Asperud 2020). Although cell death by radiation is less for the partial irradiation of 50%, the cell death by immune reaction is larger and longer than full irradiation. Measurement was available up to 30 days in the original study. (C) The growth of tumor volumes are extrapolated up to the range of 150 days.

The tumor volume at the time of treatment can be critical considering the tumor control condition described in **Eq**. (9) and as shown in **Fig. 5**. As tumor grows close to its critical volume, there is relatively small time window for successful treatment intervention. Delay in the treatment or not enough dose before the tumor outgrows the critical volume described in **Eq**. (9) may lead failure in tumor control. For this simulation, tumor is assumed left to grow for 5 extra days from the previous experiment in **Fig. 4**. Tumor volume at the time of treatment is now 0.587 cc instead of 0.250 cc in **Fig. 4**. When dose is not sufficient (10 Gy) in partial irradiation, the tumor can grow over the critical tumor volume of 1.4 cc calculated from **Eq**. (9) before immune reaction is fully developed (**Fig. 5 (A)**). On the other hand, full irradiation can control the tumor volume within the critical tumor volume with the same dose of 10 Gy, because relatively rapid volume reduction can occur in early phase. As a result, conventional full irradiation appears less sensitive to the dose prescription in this case. Therefore, special care should be taken if partial irradiation is considered when the tumor volume is approaching the critical volume. Any delay of the treatment could be detrimental. Time delay of tumor re-growth appears still larger with partial irradiation.

**Figure 5:**
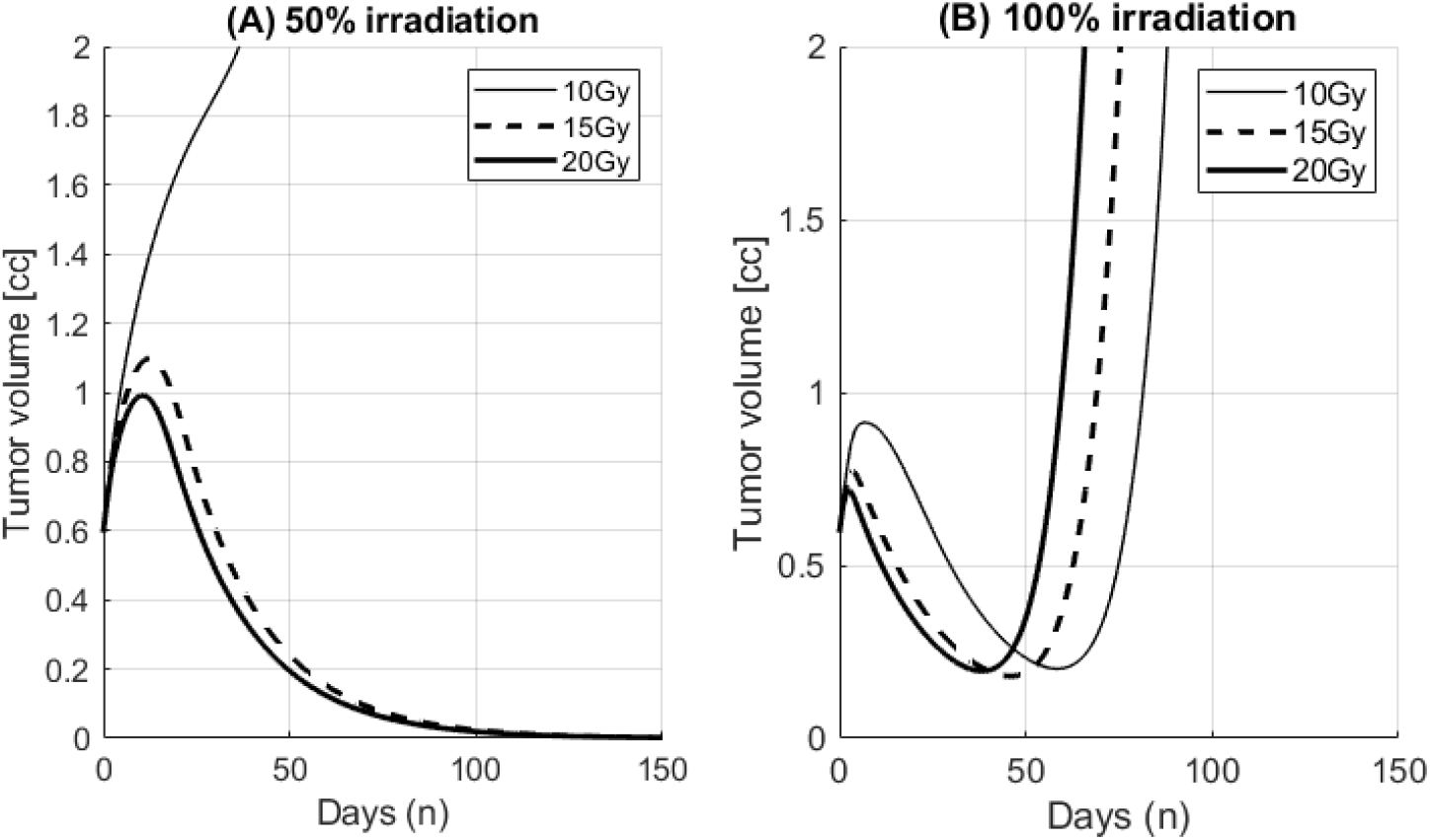
Effect of the tumor volume at treatment (critical tumor volume). In this simulation, tumor was left to grow for 5 extra days before radiation compared to the previous experiment in **Fig. 4**. Tumor volume at the time of treatment becomes 0.587 cc compared to 0.25 cc. **(A)** For the partial irradiation, dose prescription is critical for the success of treatment. When dose is not sufficient (10 Gy), the tumor volume may exceed the critical tumor volume of 1.4 cc as described in **Eq**. (9) before immune reaction is fully developed. As a result, tumors can keep growing. **(B)** On the other hand, conventional full irradiation can control the tumor volume with the same dose of 10 Gy. Relatively rapid volume reduction can hold the tumor volume below the critical tumor volume. As a result, conventional full irradiation appears less sensitive to the dose prescription parameters.

The model is applied to the tumor volume measurement of a sarcoma patient who received radiotherapy three times over 1200 days; 5 daily fractions of 6 Gy starting on day 12 after the first simulation, another 5 fractions of 6 Gy on day 609 and 5 fractions of 4 Gy on day 1148. The first fraction of the last treatment session was simultaneous integrated boost of SFRT (15 Gy at max). The immune effect triggered by radiation is shown in **Fig. 6 (A)** and the tumor volume over the course of treatments are shown in lines with the volume measurement from each CT simulation in circled marks in **Fig. 6(B) and (C)**.

**Figure 6:**
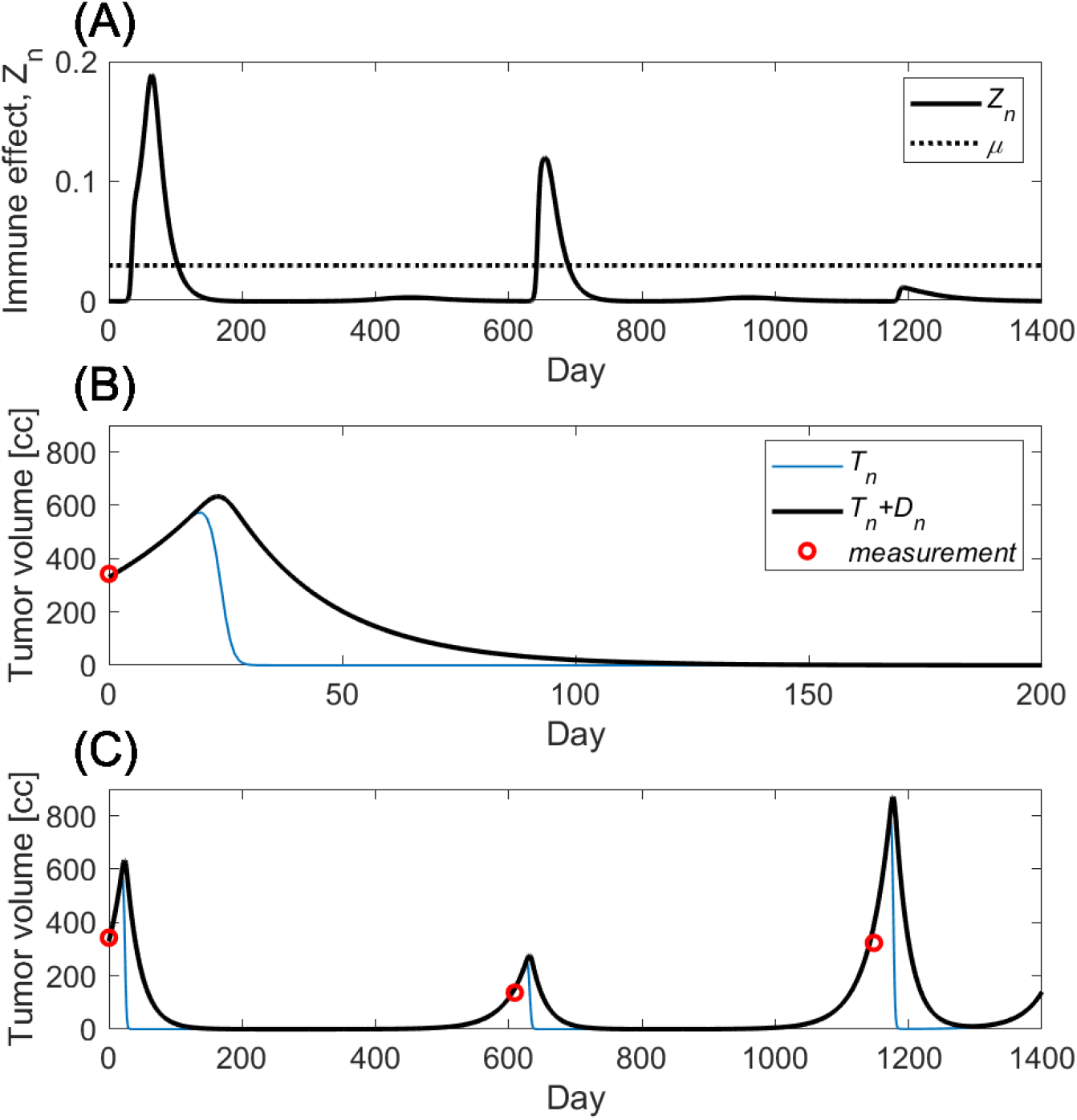
Tumor volume change in a sarcoma patient who received retreatment of radiotherapy. The patient was treated three times over 1200 days with the prescriptions of 5 daily fractions of 6 Gy (The treatment started on day 12), another 5 fractions of 6 Gy (day 609) and 5 fractions of 4 Gy (day 1148). The first fraction of the last treatment session was simultaneous integrated boost of SFRT (15 Gy at max). **(A)** shows the immune effect triggered by radiation in a solid line and growth rate *μ* in a dotted horizontal line. **(B)** shows the tumor volume change during the first session of the treatment. The tumor volume (CTV) measured from CT simulation is shown in a circled marker. **(C)** shows the tumor volume from the model (lines) and the tumor volume measurement from each CT scan (circles).

## 4 Discussion

Understanding the role and response of the immune system during and after radiotherapy is of critical importance as we move further into the era of immunotherapy. It is therefore crucial to develop mechanistic mathematical models of these phenomena to allow us to develop quantitative hypotheses. Deeper understanding of the mechanism may allow to efficiently design clinical studies aiming to amplify the synergistic effect of the radiation and the immunotherapies for cancers. In this study we extend and synthesize two previous models (Serre et al and Asperud et al). By reformulating the original equations demonstrated that two models are closely related and the effect of partial volume irradiation is explained well by the introduction of CTLs activation.

SFRT is a special radiotherapy technique with long history used before the introduction of conformal planning. It is known to be useful for large tumors difficult to manage with conventional homogeneous treatment planning. Although forgotten for few decades with the introduction of high precision radiotherapy such as IMRT/VMAT, the resurrection of this old technique is recently empowered by IMRT/VMAT (Amendola et al. 2020; Wu et al. 2020). Contrary to the collimator-based SFRT where the maximum heterogeneity is at few centimeters depth in tissue, IMRT/VMAT based SFRT can deposit the most heterogeneous dose at deeper part of the body where tumor is located. Despite the increasing interest, therapeutic gain of SFRT enabled by so called *bystander effect* is not yet fully understood. Serre et al. developed a model to explain biphasic relationship between the size of a tumor and its immunogenicity measured by the *bystander effect*. Markovsky et al. reported the delay of tumor growth in two groups of mice with 67NR tumor when treated with radiation; one with partial volume and the other with full volume irradiation. Radiation cell killing alone couldn’t explain the measurement data where half volume irradiation shows the same tumor control as full volume irradiation. Asperud et al. explained that the activation of CTLs is responsible for the development of immune response and full volume irradiation damages not only tumors but also the activated CTLs. As a result radiation cell killing of full volume irradiation may compensate immune response cell killing.

A new mathematical model was proposed in this study combining the strengths of two models. As shown in **Eq**. (2) in our mathematical representation, Serre’s model can be considered as a special case of Asperud’s model (either no dose or full dose). Starting from one of the simplest conditions (for cancers without immune suppression capability under no radiation) to the most complex condition (for cancers with fully capable immune suppression effect under radiation), the analysis of our model revealed that there are two distinct modes of tumor response: immune limited vs immune escape and its bifurcation condition. The model is capable to explain the development of immune response and the critical condition that immune response is becoming ineffective. Understanding of complex immune response and those important conditions is important in the successful design of SFRT study. With the CTLs activation at un-irradiated region, partial irradiation seems to have longer term tumor control at the expense of slower reduction of tumor volume in early phase of treatment as shown in **Fig. 4**. Since tumor cell killing by immune response is not effective anymore once the tumor grows beyond the critical volume (**Eq. 9**), SFRT with not enough dose may lose the time window for tumor control as shown in **Fig. 5**. In this case, tumor growth reaches the critical volume before immune reaction is fully developed. On the contrary, full irradiation shows more consistent results over large range of dose prescription. In other words, conventional homogeneous treatment seems to rely less on immune reaction. Therefore, it is crucial that deeper understanding of radio immune response is required for the successful applications of SFRT and special care should be taken when partial irradiation is applied if tumor volume is doubted to be close to the critical condition.

The model shows that cancers without immune suppression capability grow exponentially at first and slowly converge to the equilibrium, named as the terminal tumor volume. The tumor growth curves initially exponential and saturating to the equilibrium looks similar to other constraint growth models such as Gompertz model (Gompertz 1833). This represents the generality of the proposed model. The condition of equilibrium, however, does not mean the tumor is not life threatening.

The terminal tumor volume increases with immune suppression capability, *κ*. When *κ* is greater than the certain level called bifurcation threshold, the host body’s immune effect is suppressed below the tumor growth factor, *μ* and the tumors can grow indefinitely out of the terminal tumor volume. If the host body’s immune response is stronger, tumor growth can be limited and the terminal tumor volume is significantly smaller than those with weak immune response. In such case, the body may be able to tolerate the terminal tumor volume with finite size. When both immune response and immune suppression are strong, dynamics of tumor growth becomes unstable and tumor volume swings around the terminal tumor volume as a consequence of the competing immune dynamics. Due to the dynamic behavior where tumor volume is overshooting, tumors may be able to escape the immune limited condition earlier than the solution in **Eq**. (6) and **Fig. 3**. As shown in **Fig. 3**, tumors capable of early escape can be re-arrested to the immune limited condition when adequate amount of radiation dose is given as low as 1 Gy up to 10 Gy. Dose greater than 10 Gy was not successful because it eradicates not only tumor cells but also depletes CTLs responsible to the development of immune response. As shown in **Fig. 3 (E)**, immune response is depleted below, *μ* all the time. When the dose is too small (less than 1 Gy), it does not attenuate the vigorous dynamic behavior of tumor growth and allows early immune escape. Note that adequate amount of dose makes immune response is hanging around the tumor growth factor. The concept of “more dose the better response” may not be applicable in this case. On the contrary, SFRT in this case didn’t arrest the tumor to the immune limited mode. Instead boosted immune response at the time of radiation reduced tumor volume effectively but the immune response is forgotten afterward with tumor volume reduction. If not completely eradicated, tumor may grow back.

There are many limitations in this study. The secondary immune response, long term memory effect of immune response was not considered in the simulations for the sake of simplicity and lack of long term data for the validation of this parameter. With the secondary immune effect included, the oscillating behavior of tumor volume in **Fig. 3** may somewhat attenuate and the range of dose for re-arrest in **Fig. 3** could be extended. Other model parameters used in this study are mostly based on the mouse study reported in the literature. Application of the model to human patients should be considered carefully after thorough study on the model parameters. For this purpose the validation of model parameters is warranted with sufficient number of patient including long term data such as re-treatment of the same site. Although the model applied to 67NR tumor experiment in this study cannot be validated beyond 30 days after treatment due to the lack of measurement data, the extrapolation may be still useful to provide important insight on radio-immune response of the body. The insight gained from the study of model would allow to efficiently design clinical studies aiming to amplify the synergistic effect of the radiation and immune therapies.

## 5 Conclusion

In this study we present a mathematical model of tumor growth considering the immune response of the body and the counterpart immune suppression effect of the tumor during radiation therapy. Two distinctive modes of tumor status were identified and analyzed: immune limited and immune escape. Tumors in the immune limited mode can grow only up to a finite size and the tumor volume (terminal tumor volume, *T*_∞_) in this equilibrium can be analytically found. In the immune escape mode, there is a critical tumor mass for successful radiotherapy and it would be beneficial to start treatment early with enough dose before tumors grow more than the critical mass. The mechanism of SFRT was explained well by the development of immune response using the proposed mathematical model. Although the elevated immune response from SFRT often compensates less radiation damage in unirradiated region and produces longer tumor control effect, special attention should be paid when SFRT is considered especially tumor volume is approaching to the critical mass (**Eq. 9**). Therefore, deeper understanding of immune process would be a key for the successful application of the new cancer treatment. This study may provide some insight on the role of immune response for cancer patients by the analysis on the tumor model.

## Acknowledgements

YBC appreciates Hakan Nordstrom and Brankica Andelic from Elekta for their interest in this project and motivation on biological modeling of SFRT. JGS thanks the National Institutes of Health for their support through (5R37CA244613-02 and U54CA274513) and the American Cancer Society Research Scholar Grant (RSG-20-096-01).

